# Mapping Genetic Risk Associations to Cellular Contexts via Deep Learning and Biological Ontologies

**DOI:** 10.64898/2026.05.25.726449

**Authors:** Thomy Margalit, Hagai Levi, Ron Shamir, Ran Elkon

## Abstract

Translating genome-wide association studies (GWAS) signals into trait-relevant cellular contexts remains challenging due to the complexity of the genomic regulatory code and linkage disequilibrium among associated variants. We present a novel computational framework that aggregates deep learning–based predictions of the functional effects of noncoding variants on transcriptional regulatory elements across GWAS loci and empirically evaluates their statistical significance. By organizing these aggregated signals within biological ontologies, our approach enables statistically calibrated interpretation of GWAS associations, highlighting relevant cell-type and tissue contexts across human traits.

## Main Text

Genome-wide association studies (GWAS) have uncovered thousands of loci linked to a wide range of complex traits and diseases^1^. However, functional interpretation of these associations remains a major challenge. GWAS signals typically consist of multiple variants in strong linkage disequilibrium (LD), so each locus often contains a set of correlated candidates rather than a single causal variant^2^. This difficulty is exacerbated by the fact that most associated variants lie in non-coding regions, where effects are mediated through cell-type-specific regulatory elements^3^.

To decipher these non-coding signals, sequence-based foundation models have emerged as transformative tools^4–9^. By training on an array of epigenomic profiles recorded by multiple assays (e.g., DHS-seq, ATAC-seq, ChIP-seq), these models predict the impact of individual variants on thousands of chromatin and transcriptional profiles at base-pair resolution. While some of these models have successfully demonstrated their predictive power through validation against eQTLs and known disease-related variants^6–8^, they are primarily designed for variant-level annotation rather than trait-level inference and interpretation. To date, trait-level inference of cell-type relevance from GWAS signals has relied mainly on partitioned heritability frameworks^8,10^, most notably stratified LD score regression (sLDSC)^11^, which quantifies the enrichment of predefined genomic annotations for trait heritability.

Inspired by sLDSC, yet aiming for a more direct approach to leveraging the power of foundation models for predicting the effects of noncoding regulatory variants, we developed *IMPACT-DL* (Inference of Molecular Pathogenicity Across Cell Types using Deep Learning). Building on the recently developed Sei DL framework for sequence-based prediction of cell-type-specific regulatory effects of noncoding variants^8^, our approach performs empirical statistical tests on aggregated predicted effects across numerous cell types within LD-defined GWAS clumps.

Briefly, the *IMPACT-DL* pipeline (**Fig. 1; Methods**) begins by analyzing of GWAS summary statistics to identify lead variants and define their associated LD clumps. To ensure statistical rigor, for each clump observed in the real data, we sampled 300 mock clumps from the ethnicity-corresponding European LD reference panel of the 1000 Genomes Project^12^. These background sets were matched to the observed clumps based on local LD structure, chromosome, and minor allele frequency (MAF). We then leveraged the *Sei*^8^ model to predict the effect of every variant within the real and mock clumps on 21,907 regulatory chromatin profiles (‘22k profiles’). This collection of epigenomic profiles integrates results of DHS-seq, ATAC-seq, and ChIP-seq assays measured in 1,697 distinct cell types and tissues (**Tab S1**). To summarize these high-dimensional outputs at the locus level, we took the maximum predicted variant effect for each chromatin profile, assuming a single causal SNP per clump. Then, we averaged the top 50 clump-level scores to get a GWAS-level statistic per profile, and computed its empirical P value by comparing it to the distribution of the averaged clump-level top 50 scores from matched mock datasets. This calibration effectively isolates the regulatory signal from the confounding effects of local genomic architecture. Finally, to assess the association between a given GWAS and a specific tissue or cell type, we partitioned the 22k profiles into two groups - those corresponding to the tested tissue or cell type and all others - and contrasted the GWAS-level scores between these groups (**Fig. 1**).

**Figure 1.**
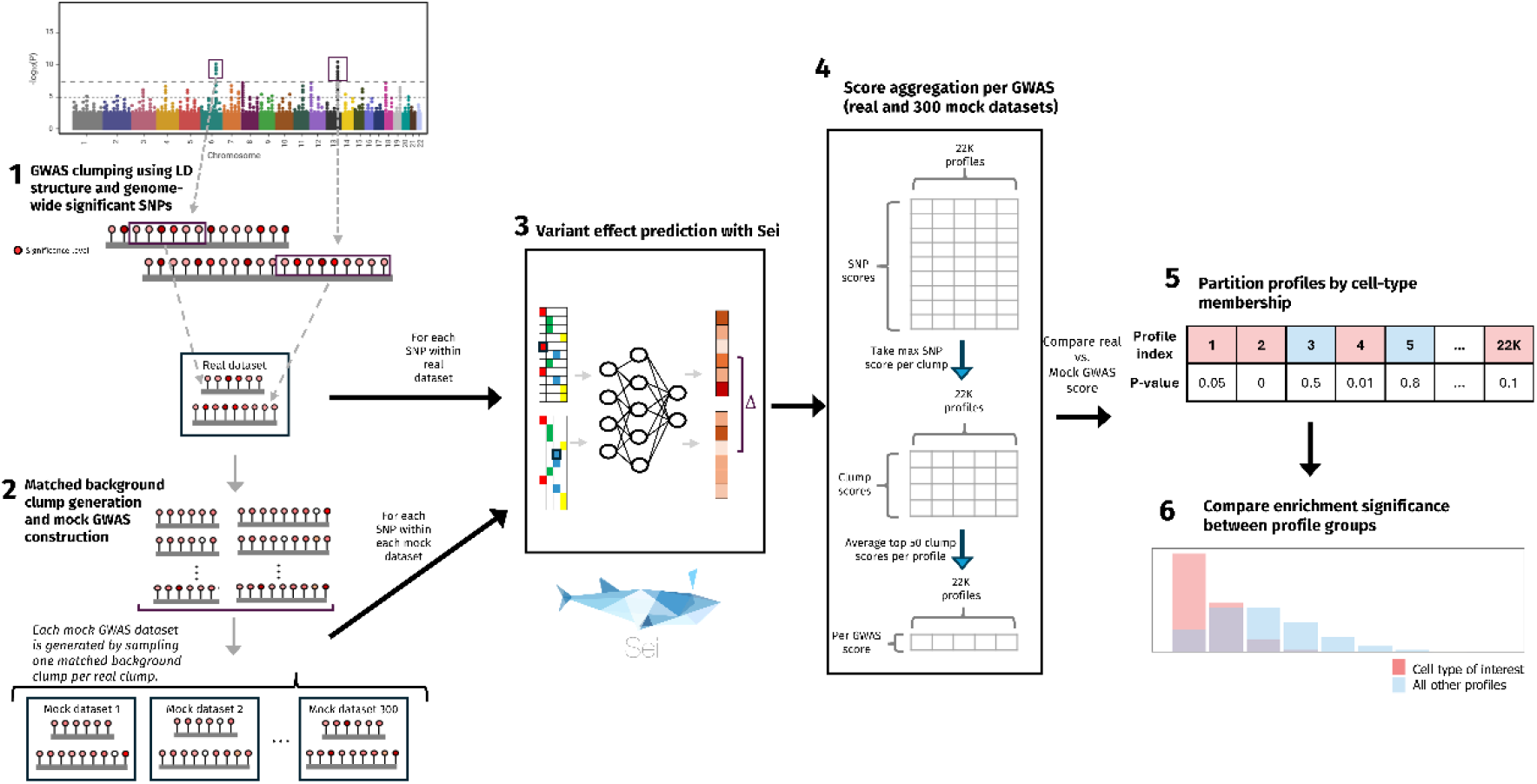
Overview of the IMPACT-DL pipeline for identifying cell types and tissues involved in the pathogenesis of human complex diseases. See Methods for a detailed description of each step.

To evaluate our method’s ability to recover known tissue associations for well-characterized traits, we applied *IMPACT-DL* to GWASs of breast cancer (BRCA), low-density lipoprotein (LDL) cholesterol levels, and atrial fibrillation (AFB), which are expected to be enriched, respectively, in cells of the mammary gland, liver, and heart. For each phenotype, we defined a subset of the 22k profiles corresponding to the relevant tissue using keyword-based annotation (see Methods) and then assessed its enrichment for signals from the three GWASs. These analyses robustly recapitulated known tissue-trait links, demonstrating the ability of our framework to capture trait-relevant regulatory programs and to pinpoint cell types involved in the molecular pathogenesis of complex traits (**Fig. 2**).

**Figure 2.**
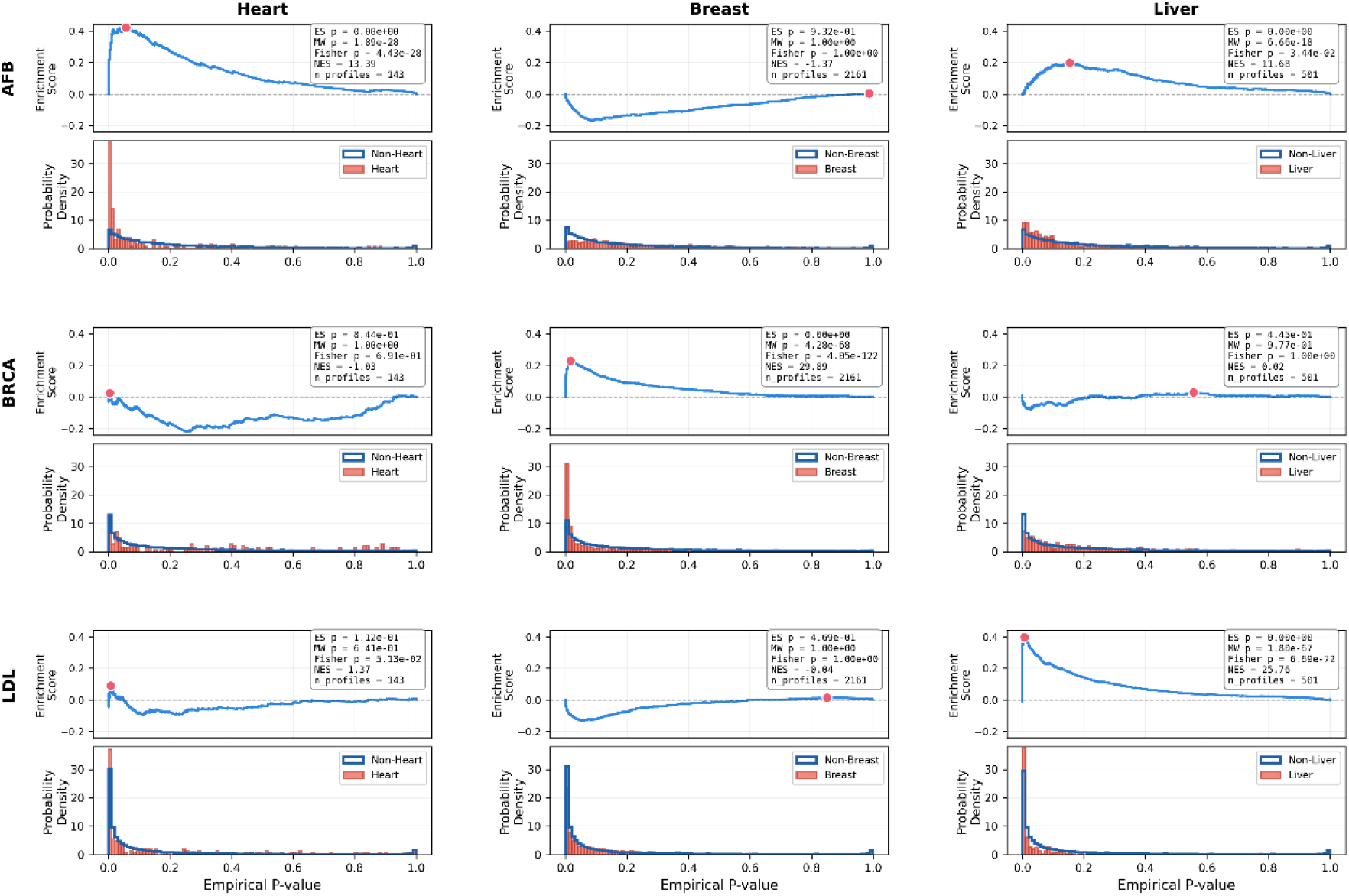
Benchmarking *IMPACT-DL* using well-characterized traits. Profile-level enrichment analysis for atrial fibrillation (AFB), breast cancer (BRCA), and LDL cholesterol (LDL) GWAS signals using manually defined sets of Sei chromatin profiles associated with the heart, breast, and liver. For each GWAS dataset-profile set pair, the top panel shows the GSEA-like enrichment curve^19^ (the red dot indicates the maximum positive enrichment score (ES), reflecting one-sided enrichment at the top of the ranked list; negative NES values indicate that the observed enrichment is weaker than expected under the permutation null), and the bottom panel shows the corresponding distributions of empirical p-values for Sei chromatin profiles inside and outside the set. P values for the enrichment of GWAS signals in the tested profile set were calculated using a permutation-based test on the enrichment score (ES), along with complementary Mann–Whitney U and Fisher’s exact tests (see Methods). (In each panel, N indicates the number of Sei chromatin profiles in the tested set).

Next, we sought to utilize *IMPACT-DL* to discover novel associations between traits and cell types/tissues. This required moving beyond manual curation of profile sets to a more systematic approach. To this end, we used the Experimental Factor Ontology (EFO)^13^, which provides a structured set of terms describing, among others, cell types and tissues, and links widely used cell lines to these terms (e.g., GM12878 → lymphoblastoid cell line → B cell → immune system). We implemented a framework that maps Sei chromatin profiles to EFO terms. By leveraging the hierarchical structure of the ontology, this approach enables grouping of chromatin profiles based on shared cellular context, even when that context is not explicitly stated in the profile label. This strategy allowed us to perform a systematic, assumption-free exploration of tissues and cell types associated with human traits and diseases based on enrichment of GWAS signals.

We first applied this automated approach to the breast cancer (BC) GWAS. Notably, we observed that regulatory noncoding signals for this GWAS were not only enriched in breast tissue and mammary epithelial cell profiles, but were also significantly enriched in a distinct branch of the EFO ontology corresponding to the lymphocyte lineage (**Fig. S1**). This finding suggests a novel link between regulatory programs active in B and T cells and genetic risk for BC. Supporting this observation, a recently published multi-ancestry BC GWAS reported enrichment of risk signals in secretory epithelial cells and innate immune cells^14^. In addition, our results highlight a shared genetic risk signal between breast and prostate cancers (**Fig. S1**), both of which are hormone-dependent cancers derived from epithelial cells of glandular tissues^15^.

Last, we performed a global analysis of 33 GWAS datasets that collectively span a broad range of phenotypes and etiologies (**Tab S2**). Hierarchical clustering of trait-term enrichments organized the results into distinct physiological axes, segregating metabolic, immune, and oncologic traits based on their regulatory signatures (**Fig. 3**; **Tab S3**). This global map revealed that while some traits exhibit highly localized signals - such as the specific enrichment of lipid traits in hepatic cellular contexts or corneal thickness in fibroblast programs, others demonstrate broader pleiotropy across diverse cell lineages. For instance, immune-mediated diseases showed distributed enrichment across the hematopoietic hierarchy, while oncologic traits captured both lineage-specific gland signatures and shared epithelial programs. For complex neuropsychiatric traits, the framework primarily identified immune signals; while these may reflect underlying neuro-immune contributions, they also highlight the current resolution limits of cell-line-derived chromatin models in specialized tissues like the brain. We repeated the analysis using three additional ontologies (Cell Ontology (CL)^16^, Cell Line Ontology (CLO)^17^, and BRENDA Tissue Ontology (BTO)^18^) and obtained highly concordant results (**Fig. S3-5**; **Tab S4-6**). Together, these results demonstrate that *IMPACT-DL* recovers established physiological associations and points to putative novel links while providing a systematic view of overlapping regulatory signatures.

**Figure 3.**
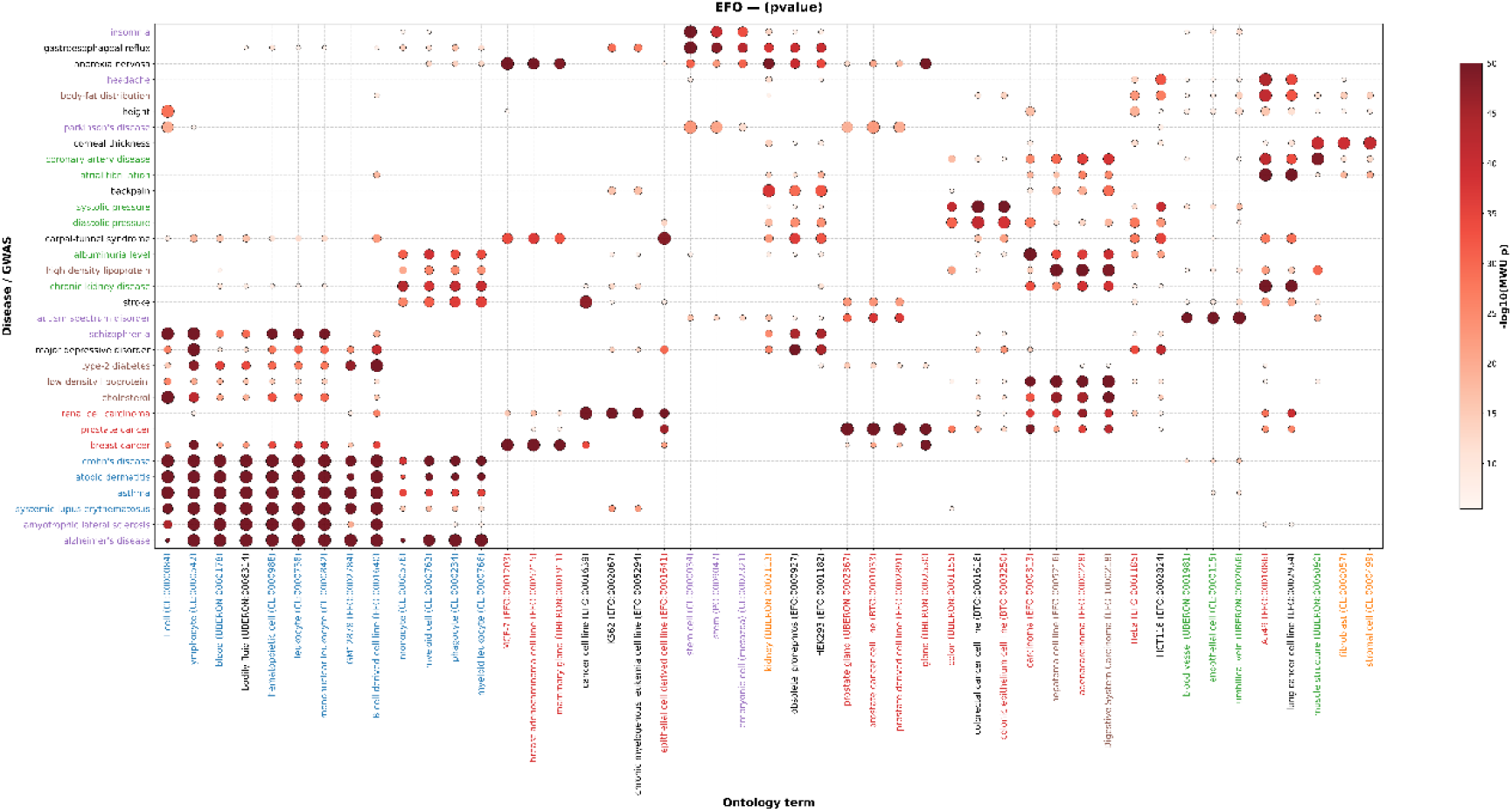
Cross-trait GWAS enrichments over ontology-defined cell type/tissue regulatory programs. Bubble heatmap showing enrichment of EFO terms (columns) across multiple GWAS traits (rows). For each GWAS, EFO terms were selected automatically by retaining the top three Bonferroni-significant terms per trait, ranked by the smallest Mann-Whitney U p-values. The union of these terms across traits is shown in the heatmap. Bubble color indicates enrichment strength (−log_10_ *MWU pvalue*). Bubble size was computed by row-wise min-max normalization of the displayed −log_10_ *MWU pvalue* values within each GWAS, so sizes indicate relative signal within a trait rather than absolute differences across traits. For visualization, −log_10_(p-values) were truncated at 50 to prevent domination by extreme signals. To reduce redundancy, ontology terms in an ancestral relationship that shared ≥80% of their associated regulatory profiles were consolidated by retaining the more specific term. Rows and columns were ordered by hierarchical clustering (average linkage). Row and column labels were color-coded according to pre-defined biological groupings assigned prior to visualization (rows: immune-blue, metabolic-brown, neuro-purple, cardio/renal-green, cancer-red; columns: liver/metabolic-brown, epithelial/oncologic-red, vascular/endothelial-green, immune/hematopoietic-blue, stromal/renal-orange, stem/neuro-purple with black indicating the “other” category).

Our results show that variant-level predictions from sequence-based foundation models can be extended to trait-level physiological interpretation by coupling LD-aware aggregation with ontology-guided organization of cellular systems, yielding interpretable maps of trait-relevant cellular contexts. This framework complements heritability-enrichment approaches such as *sLDSC*: whereas sLDSC quantifies the contribution of predefined genomic annotations to trait variance, *IMPACT-DL* directly links predicted regulatory perturbations from GWAS variants to the epigenomic landscapes of cell types relevant to disease pathogenesis.

Several limitations warrant consideration. First, our inference is constrained by the distribution and quality of available Sei’s regulatory chromatin profiles: many originate from immortalized cell lines and others from bulk tissues, limiting the tissue specificity resolution. Expanding models to regulatory chromatin profiles derived from single-cell analyses holds promise for improving this aspect. Second, ontology-based aggregation introduces sensitivity to how profiles are mapped to cell types. Because profile labels must be matched heuristically to ontology terms, and different ontologies emphasize different levels of granularity, ontology choice and matching technique may influence the result.

Despite these caveats, *IMPACT-DL* provides a novel strategy for turning foundation-model predictions into biologically structured hypotheses. As richer cell-type-resolved chromatin atlases and larger foundation models continue to emerge, we anticipate that this approach will help bridge the gap between GWAS associations and mechanistic insights, offering a transparent and statistically calibrated layer of interpretation for complex human traits.

## Methods

Methods are available in the online version of the paper.

## Acknowledgements

We thank David Groenewoud for his assistance in processing GWAS data files.

## Online Methods

### Data

We analyzed the summary statistics of 33 traits obtained from the GWAS Catalog. Only cohorts of European ancestry were included to minimize population stratification effects. A complete list of the GWASs used included in this work, along with trait descriptions and sample sizes, is provided in **Table S2**. All summary statistics were harmonized to the GRCh37/hg19 genome build.

### GWAS summary statistics quality control

GWAS summary statistics were reformatted into a unified schema prior to downstream analyses. Variants with low minor allele frequency (MAF ≤ 1%), low imputation quality (INFO ≤ 0.8), ambiguous strand alleles (A/T or C/G), or duplicated SNP identifiers were excluded. Minor allele frequencies were annotated using European population-matched gnomAD^20^ reference panels, and variants with missing frequency information were assigned a default MAF of 5%. The resulting filtered summary statistics were used for LD clumping and subsequent analyses.

### *IMPACT-DL* outline

*IMPACT-DL* identifies cell types and tissues that are enriched for predicted noncoding regulatory effects in analyzed GWAS data. It contains six main steps that are outlined here and depicted in **Figure 1**. (Subsequent sections below elaborate on each step):

1. GWAS summary statistics are used to identify genome-wide significant variants, and these variants are grouped into linkage disequilibrium (LD) clumps based on European LD structure. Each clump consists of a lead SNP and its correlated variants.
2. For each real LD clump, matched background clumps are generated from the GWAS SNPs and a European reference panel by sampling clumps from the same chromosome with similar LD structure and leader minor allele frequency, yielding 300 mock datasets per analyzed GWAS.
3. All variants from the real GWAS clumps and from each mock dataset are evaluated using the Sei sequence-based model, which predicts regulatory effects across 21,907 chromatin profiles for every variant.
4. Variant-level predictions are summarized to obtain a GWAS-level score per regulatory profile. Within each LD clump, the maximum variant effect is taken per profile, yielding clump-level scores. GWAS-level scores are computed by averaging the top (default: 50) clump-level scores across the genome, within each profile.
5. For each chromatin profile, the real GWAS score is compared against the distribution obtained from the 300 matched mock datasets to derive an empirical p-value for the profile. This procedure yields a vector of empirical p-values across all profiles, representing the trait-level regulatory signal.
6. To assess the association between the GWAS and a given tissue or cell type, the 22k profiles are divided into two groups - those corresponding to the tissue or cell type and all others, and the GWAS-level empirical p-values are contrasted between the two groups.

### LD reference construction and harmonization

To ensure that the LD estimates used reflect European ancestry, we restricted the 1000 Genomes Project ^12^ reference panel to individuals of European origin, and extracted the SNPs found in that cohort. For each GWAS analyzed, its variants were intersected with those in the filtered 1000 Genomes dataset, retaining only SNPs present in both resources. Using this harmonized SNP set, pairwise LD statistics were computed with PLINK^21^, generating an *r*^2^-based LD table describing linkage relationships between SNP pairs. This table was used for LD-based clumping and for constructing matched background datasets.

### LD-based clumping and variant selection

Clumping was performed on the GWAS genome-wide significant variants in the linkage table using PLINK^21^, with LD threshold of *r*^2^ = 0.69, 250 kb windows, and a p-value threshold of 5 × 10^−8^ (if less than 50 clumps were found, we relaxed the threshold to 5 × 10^−6^. For three GWASs this was still insufficient to yield 50 clumps, and the threshold was further relaxed to 5 × 10^−5^). All variants (clump leaders and their clumped SNPs) were considered variants of “interest”.

Variants of “interest” were exported to VCF format with reference alleles defined according to the hg19 reference genome, using PLINK2 with the -ref0from-fa flag, as required by Sei. Variants were annotated using Ensembl Variant Effect Predictor (VEP)^22^. To focus analyses on non-coding regulatory variation, and assuming that associated loci mainly contain a single causal variant, LD clumps containing protein-altering variants were excluded. Protein altering variants were identified as including one of the following annotations: *“transcript_ablation”,”stop_gained”,”frameshift_variant”,”stop_lost”,”start_lost”,”transcript_amplification”,”inframe_insertion”,”inframe_deletion”,”missense_variant”,”p rotein_altering_variant”*). The resulting VCF files were used as input for variant effect prediction with Sei. We ensured that all GWASs had more than 50 clumps left after the filtration process for downstream analysis.

### Background generation using matched LD clumps

To assess the significance of observed signals, we constructed empirical mock datasets matched to the LD structure of the selected GWAS variants of interest. Following LD-based clumping, the discovery GWAS is represented as a dataset of clumps, where each clump contains a lead SNP and all SNPs in high LD with it. We call this set the *discovery set* for the GWAS.

To generate an unbiased background, candidate background clumps were constructed exclusively from the GWAS SNPs, ensuring that background variants were subject to the same ascertainment and quality-control procedures as the discovery set. Using the precomputed LD table (described above in *LD reference construction and harmonization*), we considered all possible LD-based clumps by treating each GWAS SNP as a potential lead SNP and grouping variants with *r*^2^ ≥ 0.69 with it.

For each clump in the discovery set, we randomly identified 300 background clumps with similar genetic and structural properties. Clumps were matched on chromosome, minor allele frequency of the lead SNP (within ±0.1), number of SNPs in the clump (±15 SNPs), and clump length in kilobases (±20%). These matched clumps were used to construct empirical background datasets for downstream analyses. The matched clumps were sampled according to these criteria without replacement from the pool of all possible GWAS clumps. We did not control for genomic overlap of the clumps.

Mock datasets were constructed in the following manner. For each clump in the discovery set *c*_*j*_, let {*b*_*j*,1_, *b*_*j*,2_, …, *b*_*j*,300_} denote its matched background clumps. Mock dataset *i* was defined by selecting the *i*-th matched background clump for every discovery clump, i.e., {*b*_1,*i*_, *b*_2,*i*_, …, *b*_*m,i*_}, where *m* is the number of clumps in the discovery set. Thus, each mock set preserves the clump structure of the discovery set while replacing each real clump with a matched mock clump.

### Conversion of mock datasets to VCF files

For each mock dataset, we formed a set containing all SNPs belonging to its clumps. The resulting variant set was exported to VCF format using the same harmonization and reference-enforcement procedure applied to the discovery set. Variants were exported to VCF format using PLINK2 with reference alleles enforced according to the hg19 (GRCh37) reference genome.

VCF files were annotated using VEP, and variants annotated with protein-altering consequences were excluded, consistent with the filtering applied to the discovery set. The resulting non-coding VCF files were used as background input for sequence-based variant effect prediction with Sei.

### Prediction of effects of noncoding variants using Sei

Variants from the discovery set and from each of the 300 mock datasets were evaluated using the Sei framework. For each VCF file, variant effect prediction was performed using Sei’s command: `1_variant_effect_prediction.sh <VCF> hg19 <output_dir> --cuda`. Sei outputs per-variant effect scores across the 22k chromatin profiles. These scores were stored in HDF5 format as a variant-by-profile matrix, along with a corresponding row-label file mapping each row to its variant identifier.

### Score aggregation and significance calculation per Sei chromatin profile

The causal variant within an associated LD clump is generally unknown; therefore, each clump was treated as a set of candidate regulatory variants. For each Sei chromatin profile, variant-effect scores were summarized within each LD clump by taking the maximum absolute score across all SNPs in the clump. We regard the highest-scoring SNP per clump, for each Sei chromatin profile, as the most likely causal variant attributable to that locus under a single-causal-variant assumption.

To obtain a single GWAS-level score per profile, we aggregated across clumps by averaging the top 50 clump-level maxima. This procedure captures the most extreme regulatory signals across associated loci. The choice of *n* = 50 was guided empirically, with alternative values (*n* = 10, 30, 100) yielding comparable results (**Fig. S2**).

For each profile, the aggregated score from the discovery GWAS was compared to an empirical null distribution generated by applying the same aggregation procedure to each mock dataset. Because our hypothesis was directional, namely, that true GWAS loci would exhibit stronger regulatory effects than matched background loci: statistical significance was assessed using a one-sided empirical p-value, defined as the fraction of mock scores greater than or equal to the observed GWAS score (*p*_upper_).

### Profile-level enrichment framework

The aggregation procedure yields, per GWAS, a single empirical p-value for each of the *N* = 21,907 Sei chromatin profiles. These p-values are treated as the primary regulatory signal for downstream analyses. To evaluate the relevance of specific biological contexts (e.g., a certain cell type or tissue), we assessed whether a predefined set *S* of profiles (e.g., those corresponding to an ontology term or tissue group) exhibits significantly lower empirical p-values than the remaining *N* − |*S*| profiles.

We ranked all profiles by their empirical p-values in ascending order, such that the profile with the strongest GWAS signal has a rank of 1. Following a GSEA-like framework^19^, restricted to one-sided (positive) enrichment, for a given profile set *S*, we calculated a running sum statistic, *ES*(*i*), across the ranked list:

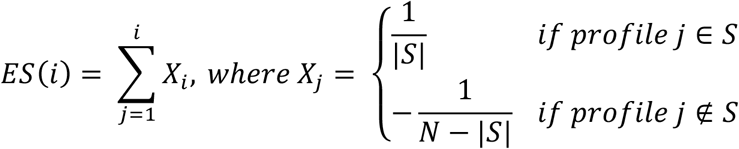

The final enrichment score (ES) is defined as the maximum value of the running sum: 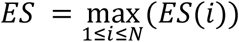. This score quantifies the degree to which profiles within the set are overrepresented at the top of the ranked list. Using permutations of the p-values, we also produce a normalized enrichment score (NES) and a p-value assessing the significance of the ES, both computed relative to the permutation-based null distribution of this ES statistic

Because empirical p-values are derived from a finite number of mock datasets (n=300), they are inherently binned, leading to frequent ties in the ranking. To ensure our results are robust to these ties, we supplement the ES with a one-sided Mann-Whitney U test. This test compares the rank distribution of the profile set against the rest, while allowing ties in ranking.

Finally, to provide a categorical measure of overrepresentation, we also calculated a Fisher’s exact test p-value. This assessed the enrichment of “significant” profiles within a set by constructing a 2*X*2 contingency table based on a p-value threshold (in our analyses, *p* = 0.1) and set membership. This approach provides a complementary, frequency-based perspective on enrichment that is independent of the overall rank distribution.

### Manual Partitioning for Initial Benchmarking

As a first validation of the framework prior to systematic ontology mapping, we partitioned the 21,907 profiles into broad tissue-specific sets using keyword matching. Profiles were assigned to a tissue if their metadata labels contained specific substrings:

- **Breast/Mammary:** “breast”, “mammary”, “MCF7”, “MCF-7”, “T47D”
- **Liver:** “liver”, “hepat”
- **Heart/Cardiac:** “heart”, “cardiac”, “cardiomyocyte”

These manually defined sets were used to generate the benchmarking results presented in Figure 2.

### Ontology matching of Sei profiles

Sei chromatin profiles were grouped into biologically defined sets using ontology-based term matching. Mapping was performed separately for four biomedical ontologies: the Experimental Factor Ontology (EFO^13^; used in the main text), and the Cell Ontology (CL)^16^, Cell Line Ontology (CLO)^17^, and BRENDA Tissue Ontology (BTO)^18^. We chose to focus on EFO for the main analysis, but all four showed comparable and interesting results, and the outputs for those analyses are provided in the supplemental material.

For each provided Sei profile label, ontology terms were identified using the BioPortal Annotator API^23^, queried separately for each ontology. To account for heterogeneity in profile naming conventions, multiple lexical variants of each label were generated, including normalization of separators (e.g., converting “B_cell”, “B-cell” to “B cell”) and removal of leading cell-line identifiers (e.g, “GM12878_B cell” → “B cell”). Non-informative or irrelevant matches were filtered out, including overly generic terms (e.g., “cell”, “tissue”), non-human organism terms, and disease-or protein-related concepts (e.g., MONDO, PR). For each Sei profile, the four top-ranked ontology matches (based on the matching score from the BioPortal Annotator API) were retained. To account for ontology hierarchy, matched terms were expanded to include their ancestor terms. Redundant nested terms covering identical sets of Sei profiles were collapsed, retaining only the most specific (leaf) terms. The resulting collapsed term-to-profile mappings defined the ontology-based sets used for downstream enrichment analyses.

### Ontology term enrichment analysis

Ontology term enrichment was performed to identify cell types and tissues whose associated Sei profiles show significant Sei variant perturbation signal for the analyzed GWAS. For each ontology term, we defined the set of Sei profiles that were mapped to that term based on the collapsed term-to-profile mappings described above. Terms represented by fewer than a minimum number of profiles (default: 10) were excluded from testing.

For a given GWAS, we used the empirical p-values derived from the Sei aggregation procedure (described in *Score aggregation and significance calculation*) as the profile-level signal. For each ontology term, we compared the distribution of *P* values for profiles associated with the term to that of all other profiles, as described in *Profile-level enrichment framework*.

To visualize the landscape of associations between GWAS risk variants and human cell types, we constructed enrichment heatmaps integrating results from multiple GWAS datasets. For each GWAS trait-ontology term pair, significance was quantified using the Mann-Whitney U p-values. To account for the high number of tests performed across ontology terms and GWASs, we applied Bonferroni correction across all comparisons, with a corrected p-threshold of 3.78e-06. This stringent strategy prioritizes the most robust biological “hotspots” and ensures that the clusters of metabolic, immune, and oncologic traits (**Fig. 3; Fig. S3-5**) represent statistically rigorous findings.

We ordered rows and columns of the heatmaps by hierarchical clustering with average linkage to help with their interpretation. The distance metrics were chosen to reflect the nature of the data: Euclidean distance was used for traits to capture absolute differences in enrichment magnitude, while correlation distance was used for ontology terms to group them by their shared enrichment patterns across traits.

## Supplementary Figures

**Supplementary Figure S1.**
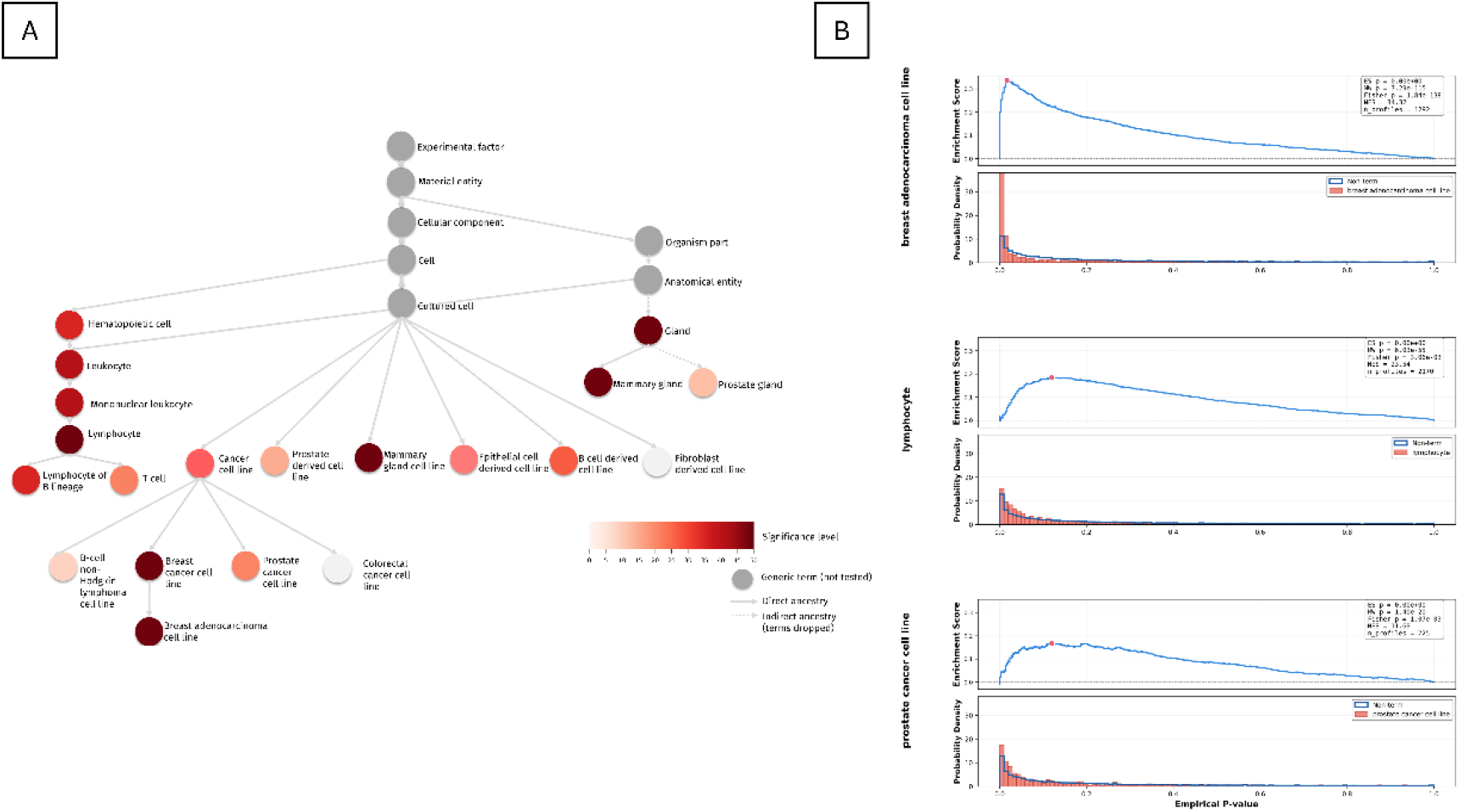
Enriched regulatory GWAS signals for breast cancer displayed on the EFO hierarchy. (a) A schematic subset of the EFO hierarchy relevant to the breast cancer GWAS, showing nodes with enriched signals and their ancestors. Nodes correspond to EFO terms, edges indicate parent-child relationships, and color intensity reflects the enrichment significance for the breast cancer GWAS. (b) Selected EFO terms enriched for breast cancer GWAS noncoding regulatory signals. For each ontology term, the top panel shows the enrichment score (ES) curve, and the bottom panel shows the distributions of empirical P values for Sei chromatin profiles within (red) and outside (blue) the term.

**Supplementary Figure S2.**
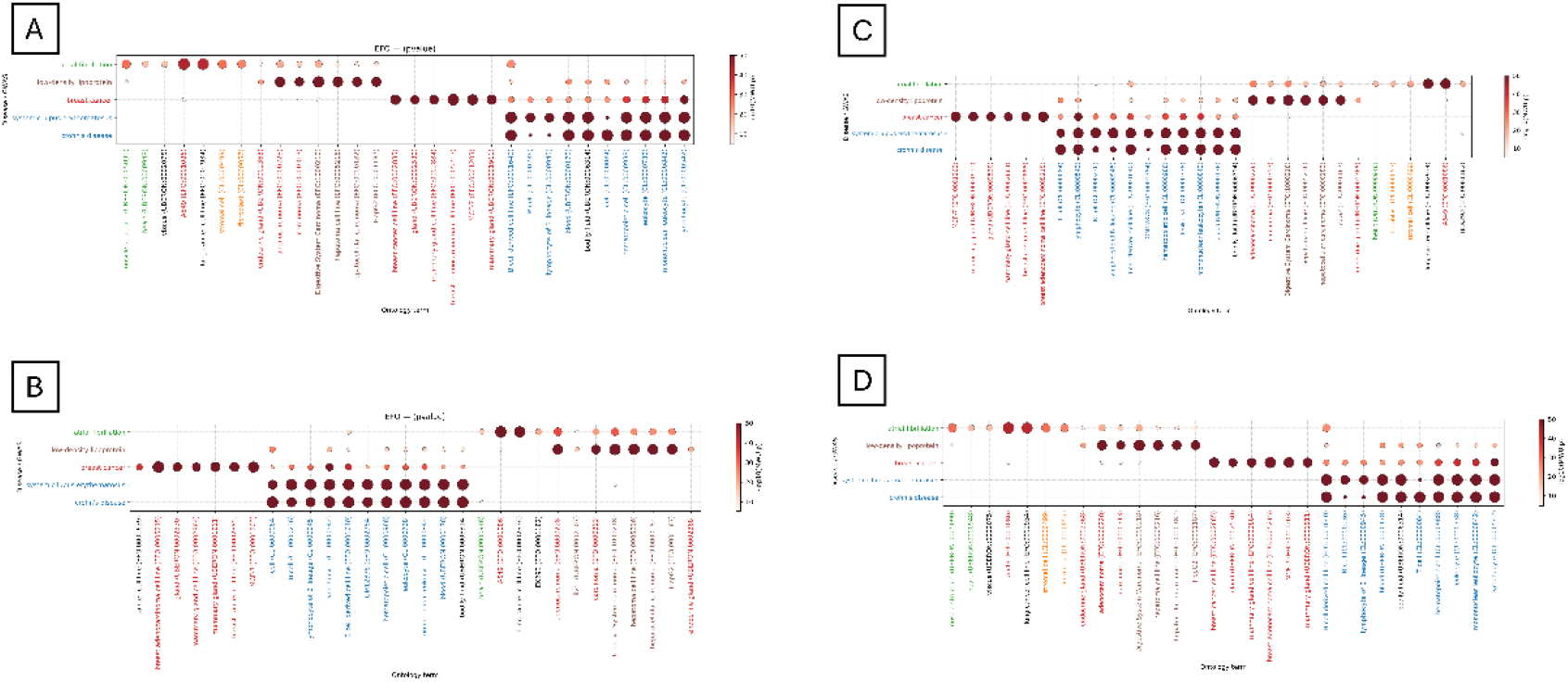
Sensitivity of GWAS-level regulatory aggregation to the number of clumps. Comparison of EFO term enrichment results obtained by aggregating the top *n* clump-level maxima per Sei chromatin profile. **(a)** *n* = 10, **(b)** *n* = 30, **(c)** *n* = 50 (default), and **(d)** *n* = 100. Five GWAS datasets were chosen, and the top 10 Bonferroni significant terms are shown per GWAS. The results were robust to the choice of *n*.

**Supplementary Figure S3.**
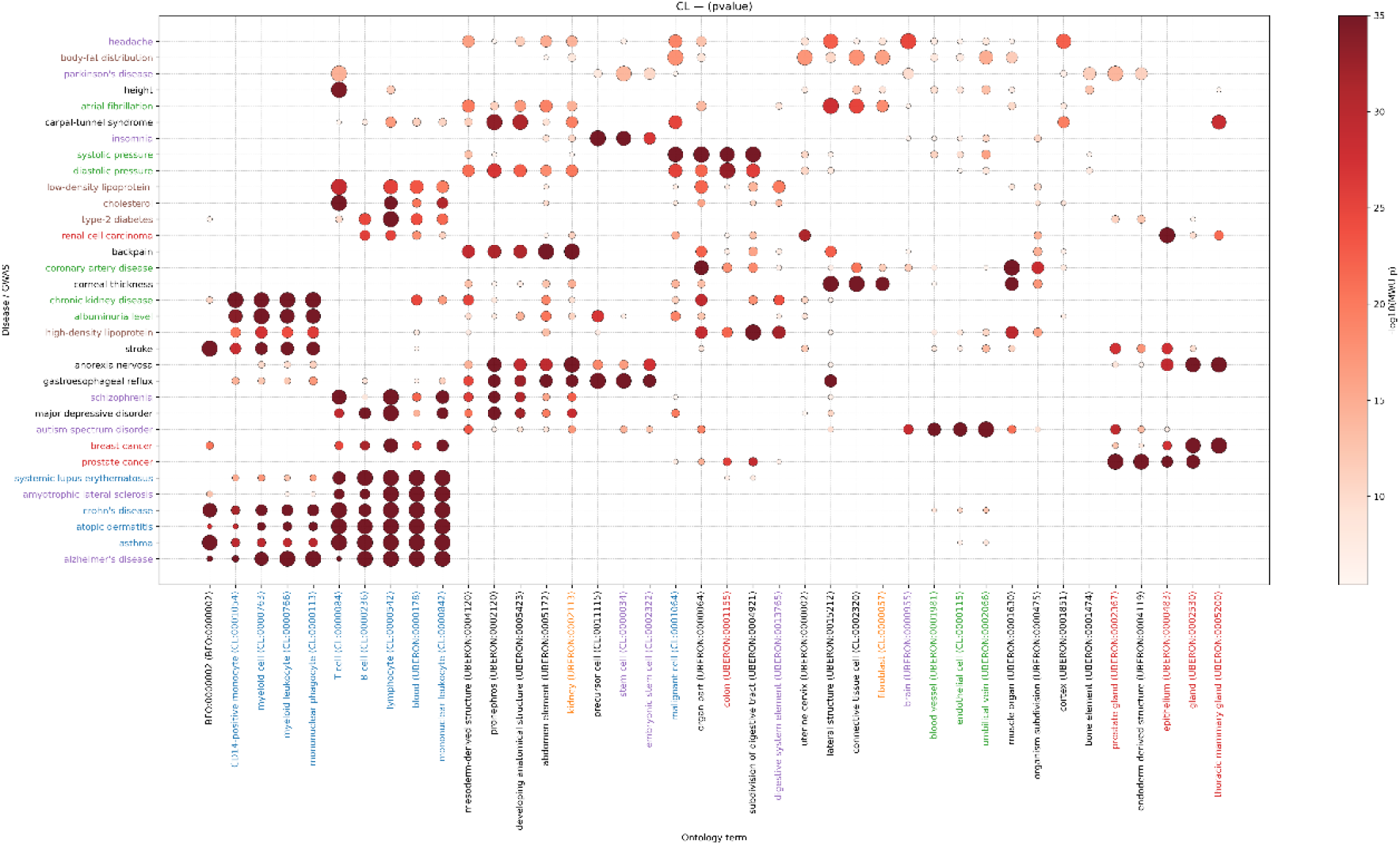
Cell Ontology-based GWAS enrichment of regulatory programs. Bubble heatmap showing enrichment of Cell Ontology (CL) terms across GWAS traits. See captions of Figure 3 for details.

**Supplementary Figure S4.**
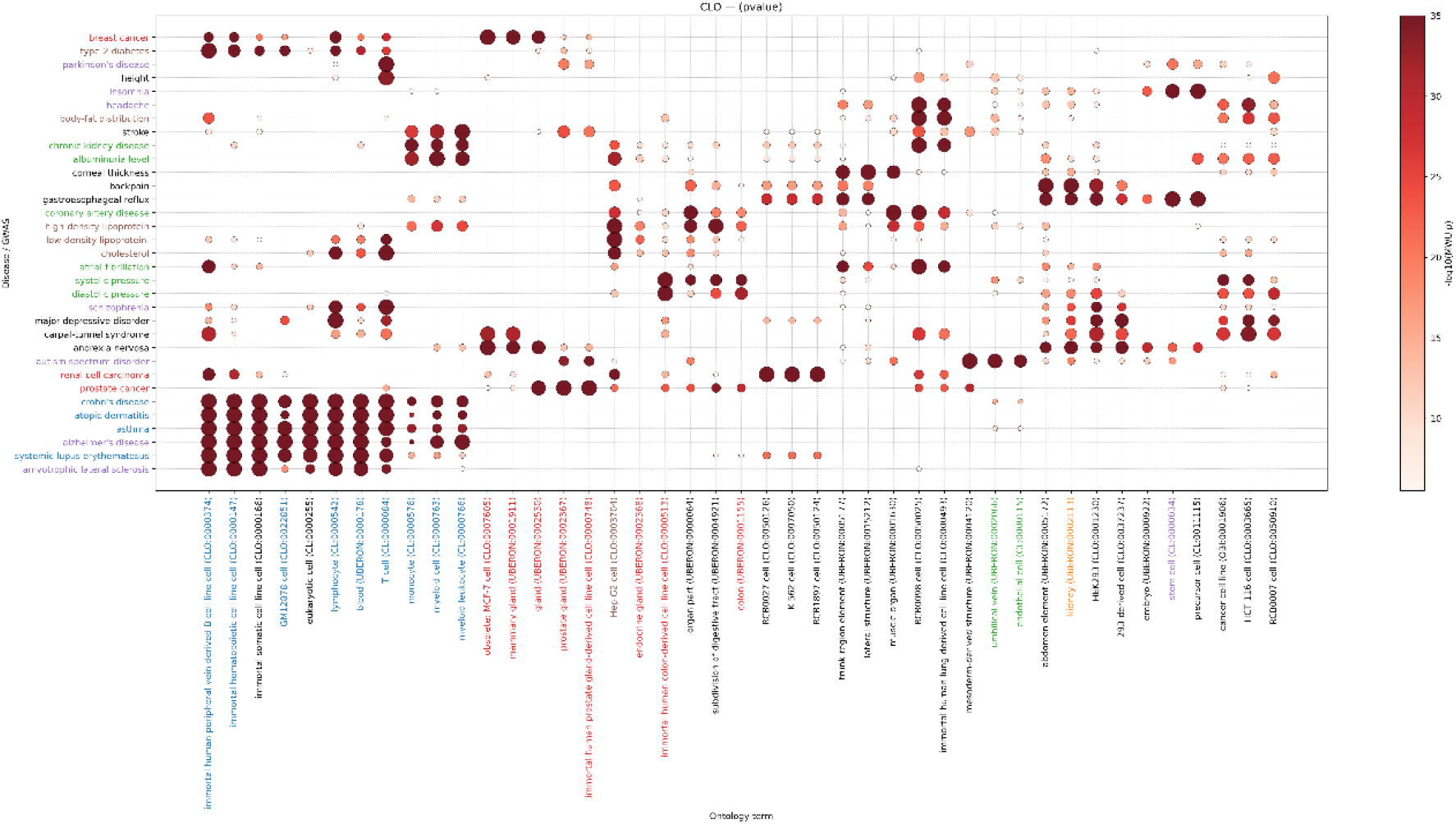
Cell Line Ontology-based GWAS enrichment of regulatory programs. Bubble heatmap showing enrichment of Cell Line Ontology (CLO) terms across GWAS traits. See captions of Figure 3 for details.

**Supplementary Figure S5.**
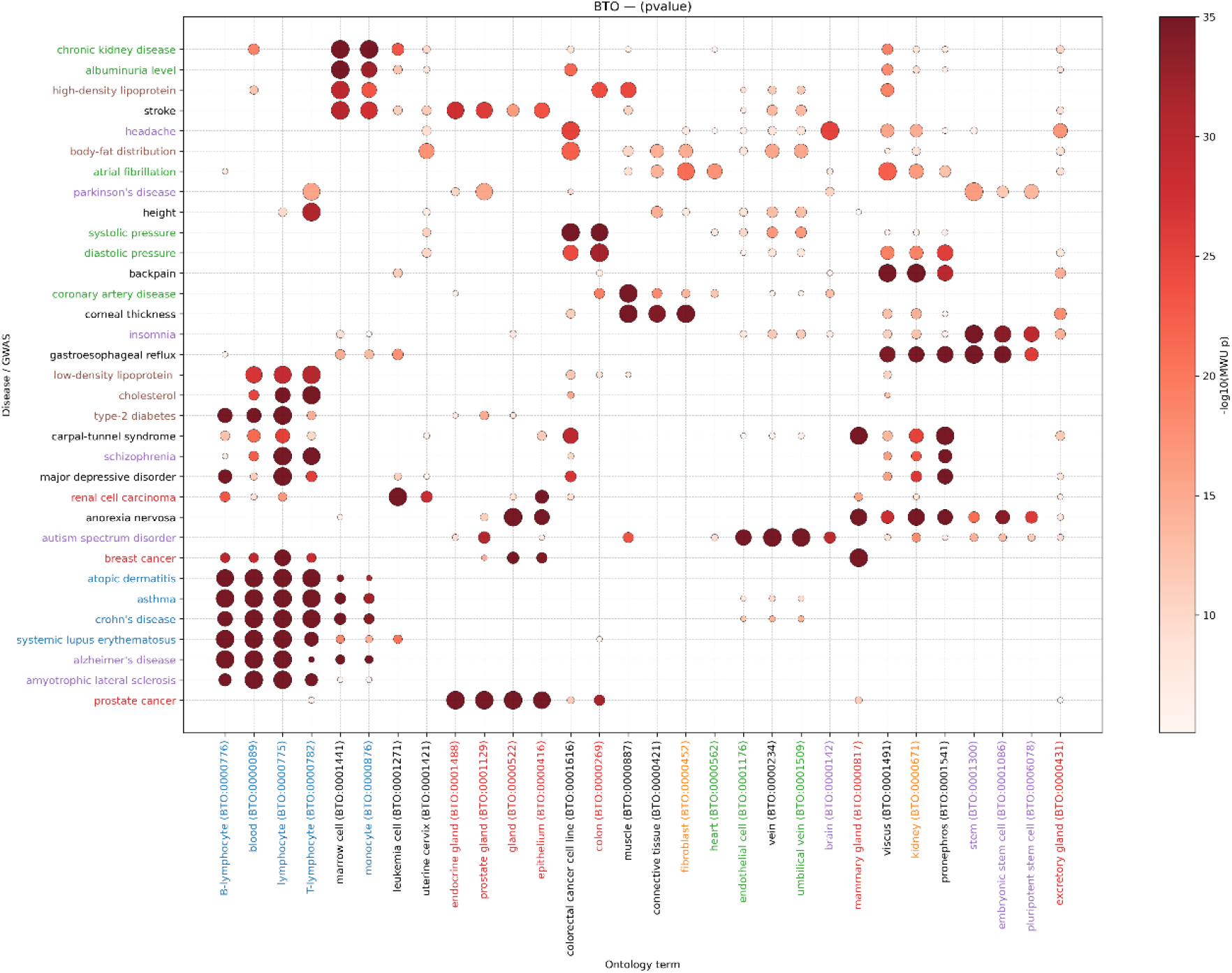
BRENDA Tissue Ontology-based GWAS enrichment of regulatory programs. Bubble heatmap showing enrichment of BRENDA Tissue Ontology (BTO) terms across GWAS traits. See captions of Figure 3 for details.**s**

## Supplementary Table Legends

**Supplementary Table 1. Sei chromatin profiles**. Excel file containing two tabs derived from the public Sei resources. **Tab 1** lists all 21,907 Sei chromatin profiles (profile identifiers and labels) used in this study, taken from the Sei public github page. **Tab 2** provides the associated biosample metadata (cell type / tissue annotations) as reported in the Sei paper (Supplementary Table 2 of the Sei publication).

**Supplementary Table 2. GWAS datasets analyzed**. Summary of GWAS datasets analyzed in this study, including trait name, source publication, sample size and ancestry.

**Supplementary Table 3. EFO-based enrichment results across GWAS traits**. Results of Experimental Factor Ontology (EFO)-based enrichment analyses for all GWAS datasets, including enrichment statistics (Mann-Whitney U p-values, enrichment scores, and multiple-testing-corrected significance values) for all tested ontology terms.

**Supplementary Table 4. Cell Ontology (CL)-based enrichment results**. Results of Cell Ontology (CL)-based enrichment analyses across all GWAS datasets. The table reports enrichment statistics (Mann-Whitney U p-values, enrichment scores, and Bonferroni-corrected significance values) for all tested CL terms.

**Supplementary Table 5. Cell Line Ontology (CLO)-based enrichment results**. Results of Cell Line Ontology (CLO)-based enrichment analyses across all GWAS datasets. The table reports enrichment statistics (Mann-Whitney U p-values, enrichment scores, and Bonferroni-corrected significance values) for all tested CLO terms.

**Supplementary Table 6. BRENDA Tissue Ontology (BTO)-based enrichment results**. Results of BRENDA Tissue Ontology (BTO)-based enrichment analyses across all GWAS datasets. The table reports enrichment statistics (Mann-Whitney U p-values, enrichment scores, and Bonferroni-corrected significance values) for all tested BTO terms.

